# Slow TCA flux implies low ATP production in tumors

**DOI:** 10.1101/2021.10.04.463108

**Authors:** Caroline R. Bartman, Yihui Shen, Won Dong Lee, Tara TeSlaa, Connor S.R. Jankowski, Lin Wang, Lifeng Yang, Asael Roichman, Vrushank Bhatt, Taijin Lan, Zhixian Hu, Xi Xing, Wenyun Lu, Jessie Yanxiang Guo, Joshua D. Rabinowitz

## Abstract

The tricarboxylic acid (TCA) cycle oxidizes carbon substrates to carbon dioxide, with the resulting high energy electrons fed into the electron transport chain to produce ATP by oxidative phosphorylation. Healthy tissues derive most of their ATP from oxidative metabolism, and the remainder from glycolysis. The corresponding balance in tumors remains unclear. Tumors upregulate aerobic glycolysis (the Warburg effect), yet they also typically require an intact TCA cycle and electron transport chain^1–6^. Recent studies have measured which nutrients contribute carbon to the tumor TCA metabolites^7,8^, but not tumor TCA flux: how fast the cycle turns. Here, we develop and validate an *in vivo* dynamic isotope tracing-mass spectrometry strategy for TCA flux quantitation, which we apply to all major mouse organs and to five tumor models. We show that, compared to the tissue of origin, tumor TCA flux is markedly suppressed. Complementary glycolytic flux measurements confirm tumor glycolysis acceleration, but the majority of tumor ATP is nevertheless made aerobically, and total tumor ATP production is suppressed compared to healthy tissues. In murine pancreatic cancer, this is accommodated by downregulation of the major energy-using pathway in the healthy exocrine pancreas, protein synthesis. Thus, instead of being hypermetabolic as commonly assumed, tumors apparently make ATP at a lower than normal rate. We propose that, as cells de-differentiate into cancer, they eschew ATP-intensive processes characteristic of the host tissue, and that the resulting suppressed ATP demand contributes to the Warburg effect and facilitates cancer growth in the nutrient-poor tumor microenvironment.

## Introduction

Animals use ATP as their main energy currency, powering functions including muscle contraction, ion pumping, and protein synthesis. ATP can be produced either from glycolysis or by mitochondrial oxidative metabolism. In the latter pathway, the tricarboxylic acid (TCA) cycle oxidizes fat and carbohydrates to make the high energy electron donors NADH and FADH2, which then drive ATP production by the electron transport chain. Catabolism of a glucose molecule in glycolysis yields two ATP, while the coupled TCA cycle and electron transport chain make around 15 ATP per TCA turn while consuming one half of a glucose molecule (or equivalently, one lactate) or one fatty-acid two-carbon unit. Accordingly, in the body as a whole, ATP production by oxidative metabolism exceeds glycolytic ATP generation by more than 20-fold.

Tumors display metabolic alterations relative to healthy tissues, likely due both to dysregulated growth signaling and to the metabolic demands of proliferation^9,10^. Pioneering studies by Warburg and others in the 1920s demonstrated that, even in the presence of oxygen, tumors rapidly convert glucose into lactate (aerobic glycolysis)^11–15^. Indeed, high glucose uptake by tumors is the basis for cancer detection by fluorodeoxyglucose-positron emission tomography^16,17^ (FDG-PET).

One potential trigger of glycolysis is impaired oxidative ATP production, due to a need to fulfill ATP demand and relief of allosteric inhibition of glycolysis. Warburg hypothesized that tumors are intrinsically respiration deficient. Initial experimental evidence was conflicting on whether tumor TCA flux was higher or lower than other tissues^11–15^. Nevertheless, the concept that tumors have defective mitochondria persisted until recent studies, which have shown that the TCA cycle and electron transport chain are important for tumor cell growth and survival, with active selection against certain mitochondrial DNA mutations in human tumors^1–6^.

Isotope tracing has been used to map fuel sources of tumor TCA metabolism. While tumors often select similar substrates to their tissue of origin, human lung cancer showed increased glucose contribution to the TCA cycle^7,8^. Comparable assessment of TCA turning rate (flux) in tumors, however, has been lacking.

TCA flux is distinct from carbon substrate selection, similar to the distinction between how fast a furnace burns fuel versus which type of fuel it takes. While the latter can be measured by steady-state isotope tracing, measuring flux requires kinetic isotope labeling measurements, as have been performed for tumor cells in culture and for selected organs, but not tumors, by NMR in vivo^18–22^.

Here, we develop and validate an isotope-tracing mass spectrometry method to measure TCA cycle flux in tissues and tumors in mice. We also quantify glucose usage flux with 2-deoxyglucose. Together, these methods show that healthy murine tissues make at least 90% of their ATP using the TCA cycle and oxidative phosphorylation. Strikingly, tumors show markedly suppressed TCA turning. Despite this decreased TCA turning, and elevated glucose flux consistent with the Warburg effect, we calculate that tumors still make a majority of their ATP oxidatively, with implied total ATP production rates in tumors significantly lower than their tissues of origin.

## Results

### Primed ^13^C-lactate infusion for TCA flux measurement

We can measure TCA flux in a tissue based on the speed of labeling of TCA metabolites and the size of the TCA metabolite pool being labeled:

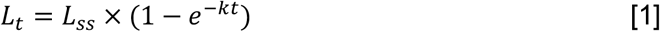

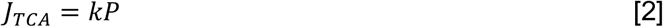

where L_t_ is the fractional labeling of tissue TCA metabolites at time t, L_ss_ is labeling of tissue TCA metabolites at steady state, k is the labeling rate constant, P is the summed concentration of TCA metabolites in the tissue, and J_TCA_ is TCA flux in a tissue (in units of nanomoles of two-carbon units consumed per minute per gram tissue)^23^. These equations apply to an idealized ^13^C-tracer that instantaneously accumulates in the bloodstream, penetrates cells, and feeds into the TCA cycle without perturbing endogenous metabolism.

Since lactate enters tissues quickly and feeds the TCA cycle in almost all tissues^7^, we hypothesized that it could potentially be used for this purpose (Figure 1A). We optimized a primed infusion of [U-^13^C] lactate (in which all three lactate carbon positions are carbon-13) so that steady-state labeling of blood lactate was attained in less than 60 seconds (Figure 1B-C). This quick approach to steady state blood labeling was important: It made sure that the rate-limiting step in tissue TCA metabolite labeling was actual turning of the cycle within the tissue (Figure 1D, Extended Data Figure 1A-B). Distinct from similar previous approaches using acetate or ethanol tracing with MRI detection^20,21^, this was achieved without markedly altering the circulating concentration of the metabolite used for tracing.

**Fig. 1.**
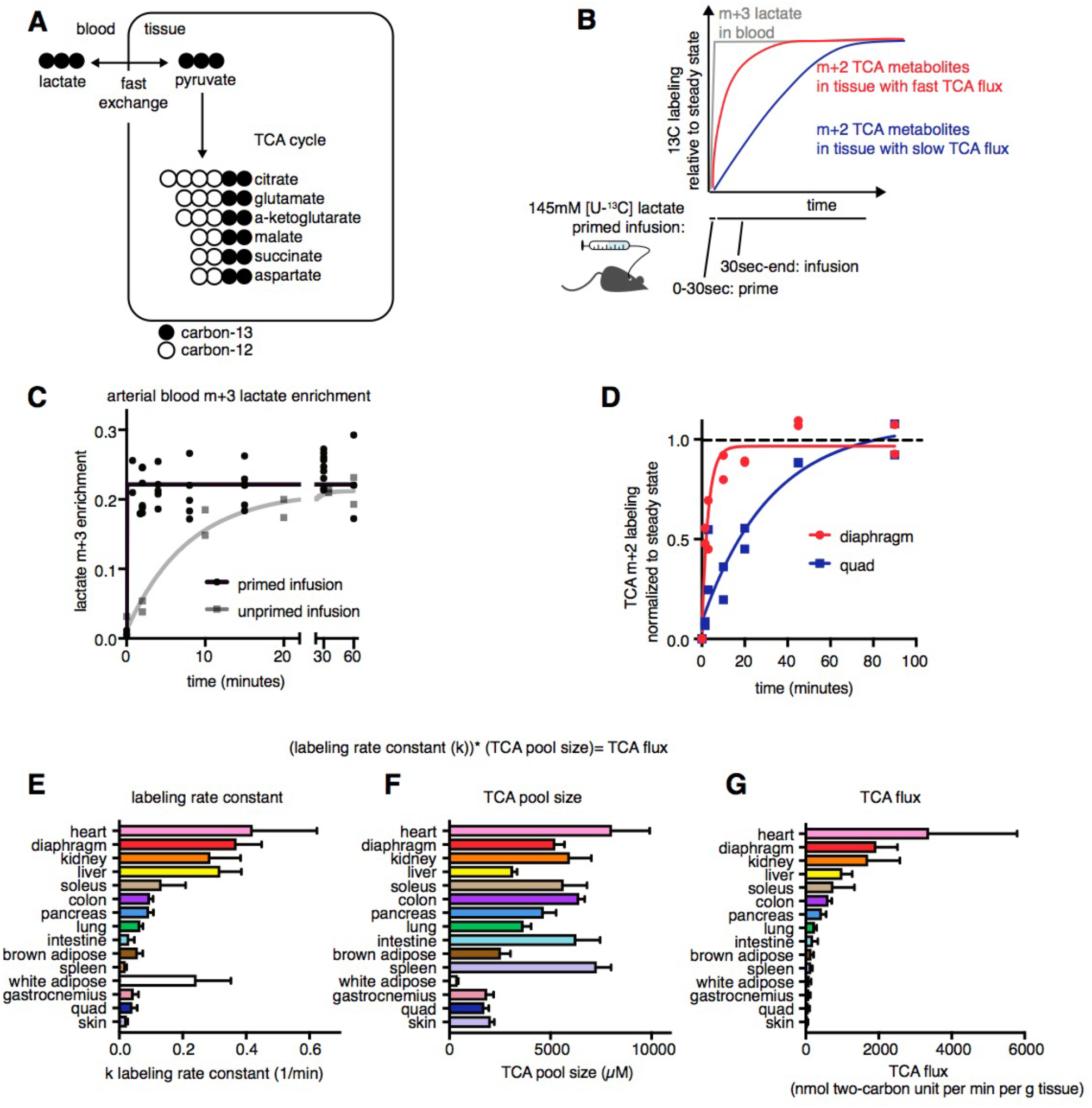
Kinetic carbon-13 lactate tracing quantifies TCA flux in vivo. **(A)** Labeling of TCA cycle metabolites from carbon-13 lactate infusion. **(B)** Schematic of labeling kinetics of blood lactate and tissue TCA metabolites from carbon-13 lactate primed infusion. **(C)** Labeling of blood lactate from carbon-13 lactate primed infusion versus non-primed infusion. **(D)** Labeling of diaphragm and quad TCA metabolites from carbon-13 lactate primed infusion. **(E)** Labeling rate constants of tissue TCA metabolites from carbon-13 lactate primed infusion. **(F)** Summed concentration of TCA metabolites in tissues (glutamate, succinate, malate, aspartate, citrate, a-ketoglutarate). **(G)** Tissue TCA fluxes. Error bars are standard deviation.

### Tissue TCA fluxes

To quantitate tissue TCA flux, we needed to measure both the labeling rate constant k and the TCA pool size P (Equation [2]). To find k, we performed carbon-13 lactate primed infusion and measured the m+2 carbon-13 labeling of tissue TCA intermediates. Due to transaminase flux being faster than TCA turning^18^, glutamate and aspartate are in labeling equilibrium with the TCA intermediates alpha-ketoglutarate and oxaloacetate respectively, and accordingly these amino acids are functionally members of the TCA carbon pool (Extended Data Figure 1C-D).

All TCA metabolites in a given tissue showed similar labeling kinetics (Extended Data Figure 1C-D), so we used the mean m+2 labeling of three well-detected metabolites, malate, succinate, and glutamate, for our calculations. We observed that different tissues accumulate TCA labeling at different rates (Figure 1D, Extended Data Figure 2A); for example, diaphragm gains TCA labeling much more quickly than quadriceps muscle. Using these labeling timepoints, we calculated the labeling rate constant k for all tissues using equation [1], with higher k representing faster labeling rate (Figure 1E).

Metabolite pool size is also required to calculate flux using kinetic measurements, because, for a given absolute flux, a larger metabolite pool takes longer to reach steady state labeling than a small pool. We used internal carbon-13 and nitrogen-15 labeled standards to quantify the tissue concentrations of TCA metabolites, including glutamate, aspartate, succinate, malate, citrate/isocitrate and succinate (Figure 1F, Extended Data Figure 2B). Oxaloacetate, succinyl-CoA, and fumarate are not well-detected by our LC-MS method, but are present at much lower abundance than aspartate and glutamate and so would not materially change the measured TCA pool sizes^18^.

By multiplying the labeling rate constant k (Figure 1E) by the TCA pool size P (Figure 1F) we calculated absolute TCA flux for each tissue (Figure 1G). We found that heart and diaphragm have the highest TCA flux per gram tissue, while skin has the lowest.

### Confirmatory TCA flux measurements with ^13^C-glutamine

Our TCA flux measurement strategy measures the speed of TCA metabolite labeling, relative to the extent of steady-state labeling from the same substrate. Accordingly, the labeling rate and measured fluxes should not depend on the tracer used. To test this concept, we turned to glutamine, a major TCA fuel which enters the TCA cycle using distinct transporters and enzymes from lactate. A disadvantage of glutamine relative to lactate is that some tissues (e.g. heart and diaphragm) use little to no glutamine in the TCA cycle, so their flux cannot be measured^25^. We developed a primed [U-^13^C] glutamine infusion strategy that achieves circulatory steady-state in less than one minute (Extended Data Figure 2C). This primed infusion was then used to measure tissue TCA labeling (Figure 2A). With the notable exception of liver, labeling rates of TCA intermediates from glutamine and from lactate were correlated (Figure 2B, R^2^ excluding liver 0.90). In liver, TCA flux measured by lactate markedly exceeded that measured by glutamine. A possible explanation is that lactate labels the TCA cycle in liver via pyruvate cycling more than TCA turning^26,27^, and thus the TCA turning rate measured with lactate is overestimated. Nevertheless, with this one exception, the agreement between rates measured with lactate and glutamine speaks to the validity of our TCA flux measurements.

**Fig. 2.**
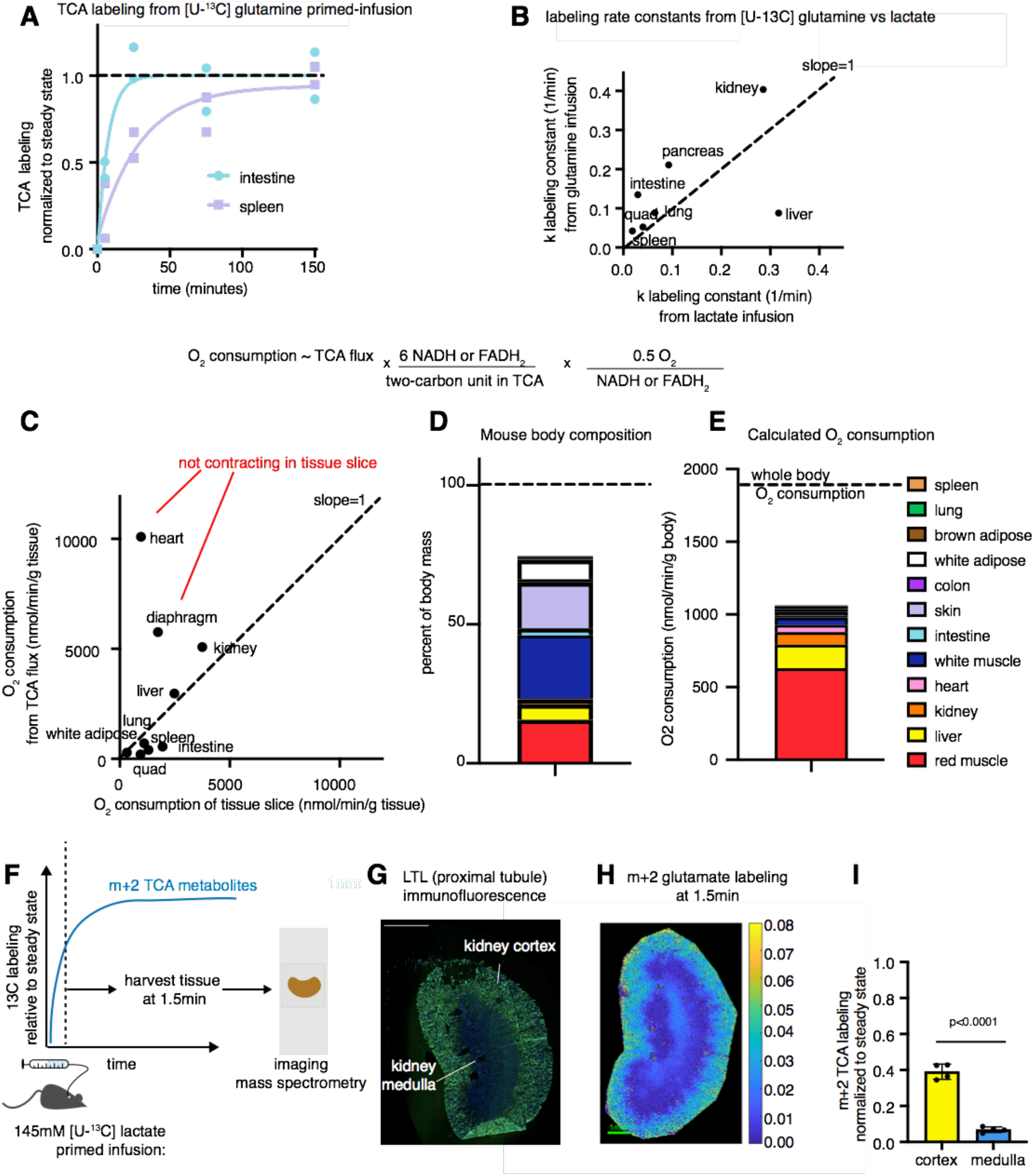
TCA flux aligns with oxygen consumption measurements. **(A)** Labeling of intestine and spleen TCA metabolites from carbon-13 glutamine primed infusion. **(B)** Labeling rate constants of TCA metabolites measured by carbon-13 glutamine versus lactate primed infusions. **(C)** Oxygen consumption calculated from carbon-13 lactate TCA flux measurement versus oxygen consumption previously measured in tissue slices. **(D)** Fraction of mouse body composed of each tissue^30^. **(E)** Calculated oxygen consumption by each tissue (based on tissue mass and TCA flux measured by carbon-13 lactate). Dotted line is whole-body oxygen consumption measured by a metabolic cage. **(F)** Schematic of spatial measurement of TCA flux using carbon-13 lactate primed infusion and imaging mass spectrometry. **(G)** Immunofluorescence staining of kidney proximal tubule using lotus tetragonolobus lectin. **(H)** Imaging mass spectrometry of glutamate m+2 labeling after 1.5 min carbon-13 lactate primed infusion. **(I)** Quantification of cortex versus medulla TCA metabolite labeling. Error bars are standard deviation, p value is from an unpaired two-tailed t test.

### Agreement with oxygen consumption measurements

To further validate our TCA flux measurement method, we compared oxygen consumption calculated from our TCA measurements to oxygen consumption directly measured in ex vivo tissue slices^28^. Due to redox constraints, oxygen consumption is directly proportional to TCA flux. Either lactate or fatty acid oxidation produces 6 reducing equivalents in the form of NADH or FADH2 per acetyl-CoA. Each reducing equivalent contributes two electrons to the electron transport chain resulting in the consumption of ½ molecule of O_2_:

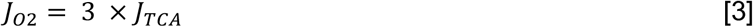

where J_O2_ is oxygen-consumption flux. We found that our measurements of TCA turning correlated well with historical tissue slice data, with the exception of heart and diaphragm (Figure 2C, R^2^=0.85 excluding heart and diaphragm). In heart and diaphragm, we observed greater TCA turning than anticipated based on *ex vivo* tissue slice oxygen consumption, likely reflecting active muscle contraction driving TCA flux *in vivo*. Consistent with this, in perfused beating mouse hearts, oxygen consumption was 20-fold greater than in tissue slices^29^, and only 2-fold different from our *in vivo* measurements.

To quantify the contribution of each tissue to whole body oxygen consumption, we multiplied the oxygen consumption calculated from tissue TCA fluxes by the fractional mass of each tissue in the body (Figure 2D-E)^30–33^. We found that red working muscles like diaphragm and soleus, which make up approximately 16% of body mass, accounted for a majority of whole-body oxygen consumption. Adding up the oxygen consumption by all tissues yielded an estimate of 1026 nanomoles O_2_ consumed per minute per gram body weight, which is on the same order of magnitude as the true measured oxygen consumption measured by a metabolic cage, 1930 nmols/min/g ^25^. The missing body oxygen consumption likely reflects unsampled tissues (e.g. bone) which comprise 30% of body mass, reactions that directly use oxygen without passing through the TCA cycle (e.g. fatty acid desaturation and peroxisomal oxidation) and/or underestimation of TCA turning rate in high flux tissues (e.g. heart, diaphragm, and kidney, where tissue lactate labeling is not faster than tissue TCA labeling, Extended Data Figure 1A-B).

### Imaging mass spectrometry of kidney TCA flux

The experiments above measured TCA flux averaged across each organ as a whole; however, tissues are composed of different cell types with distinct metabolism. For example, the kidney cortex is responsible for sodium and potassium pumping in order to recover metabolites from the glomerular filtrate, and thus has a higher energy demand than the kidney medulla^34^. To measure regional TCA flux across the kidney, we performed a primed infusion of carbon-13 lactate as above, then after 90 seconds isolated and froze the kidney, sectioned it, and performed imaging mass spectrometry. This pre-steady state TCA metabolite labeling, which correlates with flux, was much higher in the kidney cortex than the medulla (Figure 2F-I), and moreover within the cortex showed a gradient from peripheral to central, with the most superficial region showing the highest flux. We are unaware of other technologies for imaging energy production on this spatial scale.

### Kinetic 2-deoxyglucose infusion quantifies glucose usage flux *in vivo*

Two pathways generate ATP in cells, the TCA cycle (in combination with the electron transport chain) and glycolysis. Thus to estimate the routes of total tissue ATP production we also measured *in vivo* glucose usage flux. We took advantage of the glucose analog 2-deoxyglucose, which enters tissues using glucose transporters and is phosphorylated by hexokinase but then cannot be metabolized further and remains trapped in tissues^35^. Clinically, fluorodeoxyglucose positron emission tomography (FDG-PET) uses a similar strategy to measure glucose usage in tissues. As typically performed it does not yield quantitative values of glucose flux, though kinetic FDG-PET experiments can measure it^35–37^. Similarly, tritiated 2-deoxyglucose injection has been used to measure relative glucose usage across tissues^38^.

We intravenously infused mice with [1-^13^C] 2-deoxyglucose and used mass spectrometry to measure 2-deoxyglucose levels in blood and accumulation of 2-deoxyglucose-phosphate in tissues over time (Figures 3A-C). Using [1-^13^C] 2-deoxyglucose increased the sensitivity of this measurement, because tissues displayed high background at the expected mass of carbon-12 2-deoxyglucose-phosphate (Extended Data Figure 3A). The resulting data yield tissue glucose usage flux:

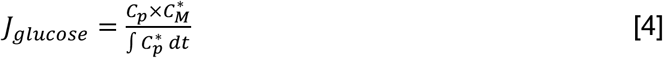

where J_glucose_ is the rate of glucose uptake and phosphorylation, C_P_ is glucose concentration in the blood (Extended Data 3C), C_M_* is 2-deoxyglucose-phosphate concentration in the tissue, and the integral of C_P_* is the integrated concentration of the blood 2-deoxyglucose concentration with respect to time up to that timepoint^35^. We found that 2-deoxyglucose infusion up to 15 minutes yielded a linear relationship between tissue 2-deoxyglucose-phosphate concentration and the integral of blood 2-deoxyglucose concentration, demonstrating that in this time frame the glucose import and phosphorylation machinery was not saturated (Figure 3C, Extended Data Figure 3B, 3D-E). Glucose flux per gram tissue was highest in brown adipose, diaphragm, brain, and soleus; lowest in skin; and correlated well with FDG-PET^39^ (Figure 3D, 3E, R^2^=0.93).

**Fig. 3.**
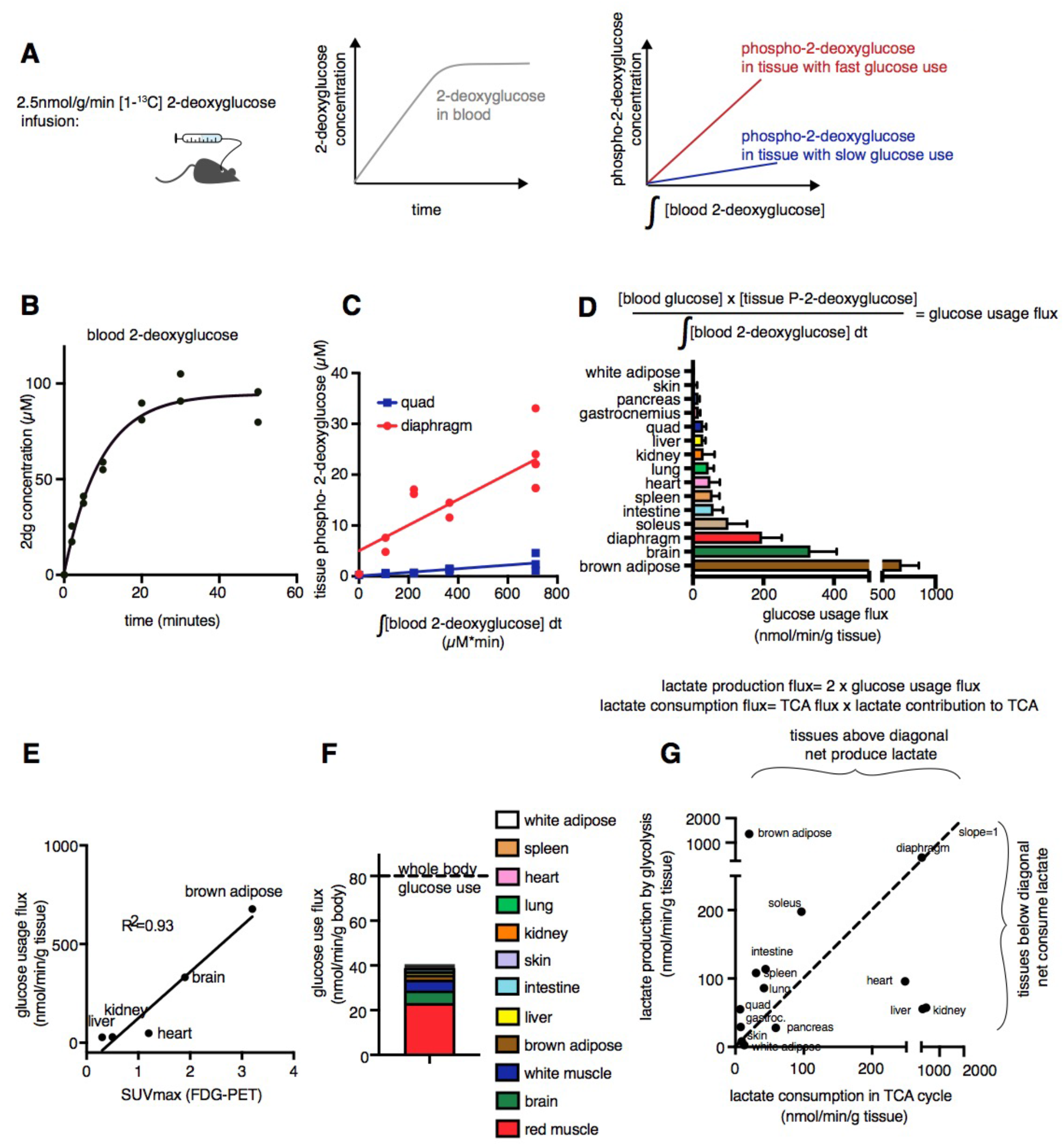
Kinetic 2-deoxyglucose infusion quantifies glucose usage flux in vivo. **(A)** Schematic of [1-^13^C] 2-deoxyglucose infusion. **(B)** Concentration of [1-^13^C] 2-deoxyglucose versus time in blood during infusion. **(C)** Concentration of diaphragm and quad [1-^13^C] 2-deoxyglucose-phosphate versus integral of blood [1-^13^C] 2-deoxglucose with respect to time, during 2-deoxyglucose infusion. **(D)** Tissue glucose usage flux. Error bars are standard deviation. **(E)** Correlation between absolute glucose usage measured by [1-^13^C] 2-deoxyglucose infusion and relative glucose usage measured by FDG-PET^39^. **(F)** Calculated glucose usage by each tissue at the whole-body level. Dotted line is whole-body glucose turnover rate measured by [U-^13^C] glucose infusion. **(G)** Production and consumption of lactate by tissues calculated from glucose flux and from TCA flux measured by carbon-13 lactate multiplied by lactate contribution to TCA cycle.

To calculate the contribution of each tissue to whole-body glucose flux, we multiplied the calculated glucose usage flux by the fraction of the body composed of each tissue^30–33^. This revealed that red working muscles like diaphragm and soleus are responsible for a majority of the body’s glucose usage (Figure 3F). Taken together with Figure 2E, this observation suggest that red working muscles are the predominant tissue type responsible for whole-body nutrient consumption. The sum of tissue glucose flux was somewhat less than whole-body glucose turnover flux (41 versus 80 nanomolar glucose/minute/gram body weight, dotted line in Figure 3F).

Classically, it is thought that cells direct carbon exiting glycolysis into the TCA cycle for oxidation. However, cells may also acquire carbohydrate for TCA burning by taking up lactate. Isotope tracing reveals extensive use of lactate to feed the TCA cycle^25,40,41^ but up to now methods to determine whether there is net lactate uptake, or merely exchange with carbon coming from glycolysis, have been limited to arterio-venous sampling. We estimated lactate production by each tissue as two lactate molecules produced per glucose used, and calculated lactate consumption in the TCA cycle as the product of TCA flux and the fraction of the TCA cycle contributed by lactate (Extended Data Figure 2D). Consistent with prior arterio-venous measurements, this approach identified kidney, liver, and heart as net lactate consumers^42,43^ (Figure 3G). Thus, our glucose usage measurements align well with prior orthogonal experimental approaches.

### Tumors have lower TCA flux than healthy tissues

Having established methods to measure TCA and glucose flux in tissues, we then applied these methods to tumors. Historically, studies have consistently found high rates of glucose usage in tumors, but (in part because tumors often lack a well-defined draining vein) there has been no reliable way to measure their TCA turning rate. To this end, we measured TCA flux in tumors using both carbon-13 lactate and carbon-13 glutamine primed infusions. We examined five tumor models: genetically engineered mouse models (GEMM) of pancreatic cancer and lung cancer, flank-implanted pancreatic- and lung-derived tumors, and a human colon cancer xenograft model (denoted respectively as GEMM PDAC, GEMM NSCLC, flank PDAC, flank NSCLC, and CRC xeno). Strikingly, all tumor models examined had much lower TCA flux than the corresponding healthy tissue: tumor TCA flux was 3.3- to 25-fold lower than corresponding healthy tissue flux depending on the tumor type (Figure 4A-P). TCA labeling from carbon-13 was gated by TCA turning, not perfusion or lactate import (Extended Data Figure 4A-B). These data suggest that tumors can satisfy their requirements for energy, biosynthesis, and redox balancing with relatively low TCA flux.

**Fig. 4.**
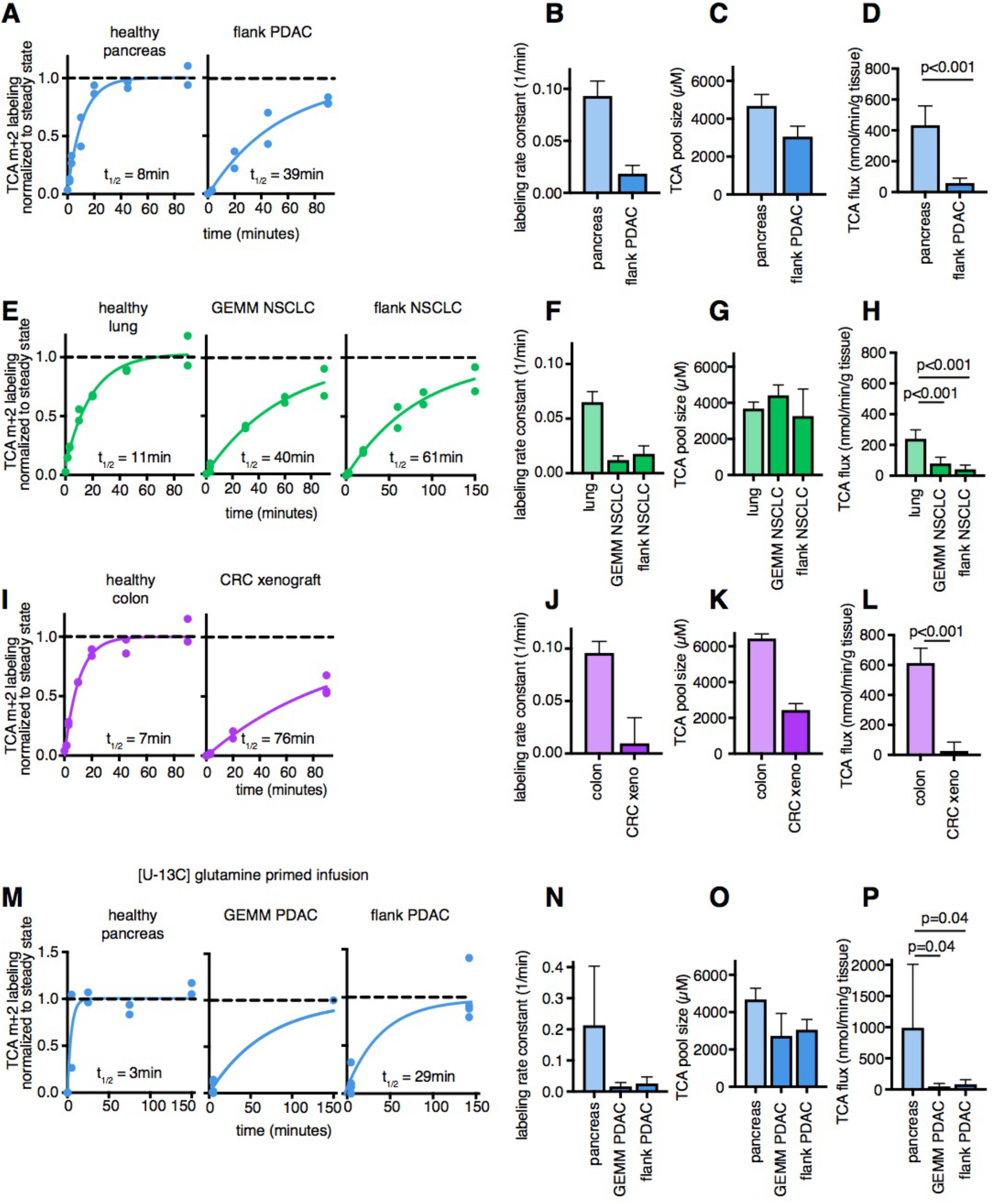
Tumors have lower TCA flux than healthy tissues. **(A)** Labeling of TCA metabolites from carbon-13 lactate primed infusion in healthy pancreas and in a flank-implanted model of pancreatic adenocarcinoma (Pdx1-cre LSL-Kras^G12D/+^ p53^R172H/+^, “flank PDAC”). **(B)** Labeling rate constants from carbon-13 lactate primed infusion, **(C)** summed concentration of TCA metabolites, and **(D)** TCA flux in pancreas and flank PDAC. **(E - H)** Analogous data for healthy lung, a genetically-engineered mouse model of non-small cell lung cancer (Adenovirus-cre LSL-Kras^G12D/+^ Stk11^-/-^p53^-/-^, “GEMM NSCLC”), and a flank-implanted model of non-small cell lung cancer (Adenovirus-cre LSL-Kras^G12D/+^ p53^-/-^, “flank NSCLC”). **(I - L)** Analogous data for healthy colon and a xenograft of the human colorectal cancer cell line HCT116 (“CRC xeno”). **(M - P)** Analogous data for healthy pancreas, a genetically-engineered mouse model of pancreatic adenocarcinoma (Pdx1-cre LSL-Kras^G12D/+^ p53^-/-^, “GEMM PDAC”), and flank PDAC, except using carbon-13 glutamine infusion. All error bars are standard deviation, p values are from a Welch’s two-tailed t test.

### Tumors have similar or higher glucose usage flux than healthy tissues

To identify how much tumor energy comes from glycolysis versus oxidative metabolism, we next measured glucose usage flux in tumors using kinetic [1-^13^C] 2-deoxyglucose infusion. In all four models examined, GEMM lung tumor, flank pancreatic and lung tumors, and colon cancer xenograft, tumors displayed somewhat higher glucose flux than the corresponding healthy tissue (Figure 5A-D). The increase in glucose usage varied depending on the tumor type, ranging from around 1.5-fold higher (flank lung tumor, not significantly different from normal lung) to 3.6-fold higher (GEMM lung tumor, p=0.007).

**Fig. 5.**
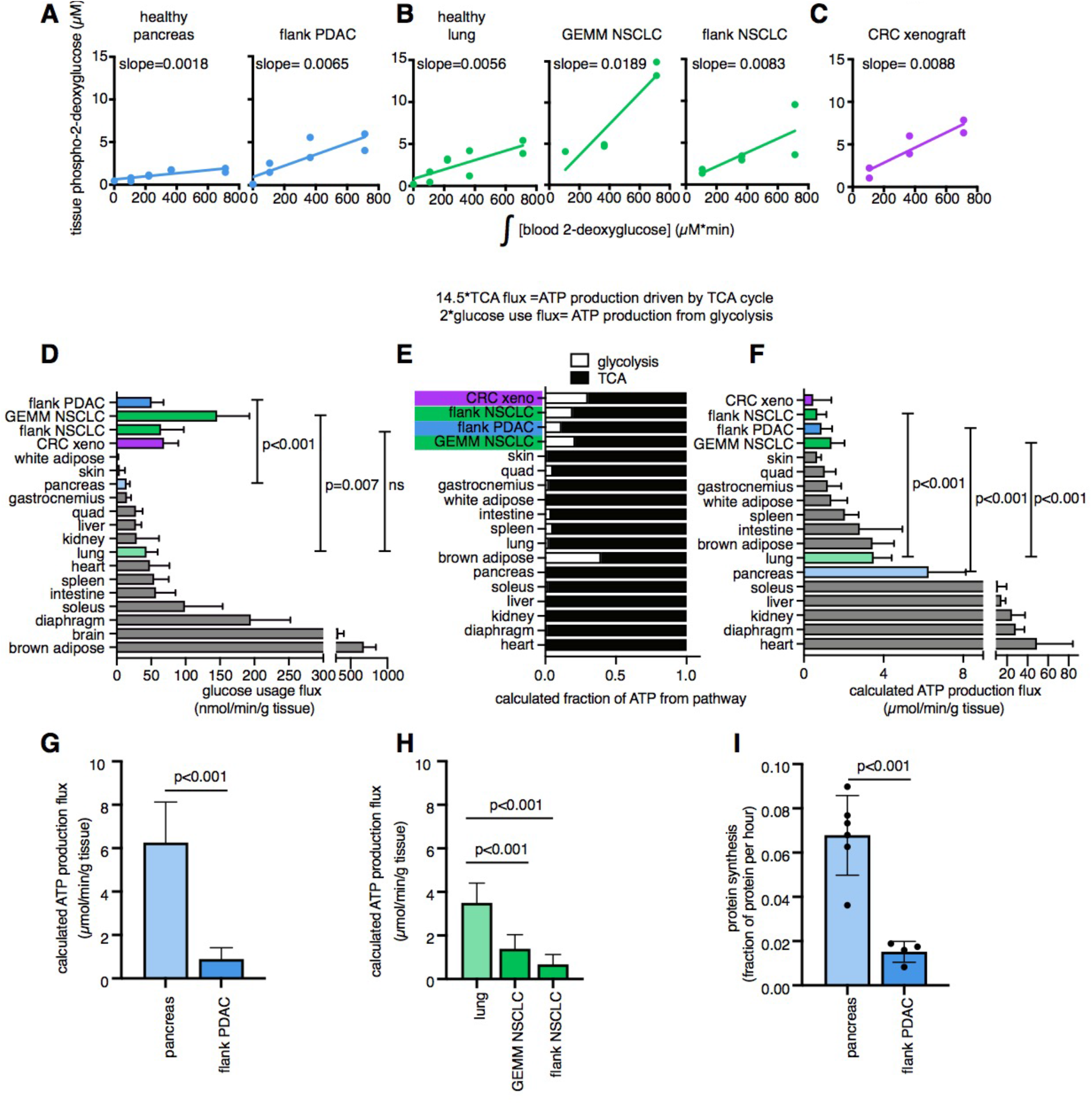
Tumors make ATP slower than healthy tissues. **(A)** Concentration of [1-^13^C] 2-deoxyglucose-phosphate versus the integral of blood [1-^13^C] 2-deoxyglucose in healthy pancreas and flank PDAC, **(B)** in healthy lung, GEMM NSCLC, and flank NSCLC, **(C)** in xeno CRC. **(D)** Glucose usage flux in healthy tissues and cancer models, ns is not significant (p=0.20). **(E)** Calculated fraction of cellular ATP derived from glycolysis or from TCA cycle coupled to electron transport chain. **(F)** Calculated total ATP production flux, t-tests compare tumors to corresponding healthy tissue. **(G)** Calculated total ATP production flux (same data as figure F), displaying pancreas and flank PDAC or **(H)** lung and lung tumors. **(I)** Protein synthesis rate of healthy pancreas and flank PDAC measured based on protein labeling from infused carbon-13 valine. All error bars are standard deviation, p values are from Welch’s two-tailed t test.

Consistent with classical Warburg metabolism, we calculate that tumors net release lactate, as they have higher estimated lactate production from glycolysis than lactate burning in the TCA cycle (7- to 53-fold higher lactate production than lactate consumption in TCA depending on tumor type, Extended Data Figure 4C). Note that tumors use circulating lactate as a TCA fuel, however they release more lactate produced from glycolysis than they oxidize in the TCA cycle.

### Tumors make ATP slower than healthy tissues

A main goal of this study was to find how much the TCA cycle contributed to energy production in tumors. Using our glucose and TCA fluxes, we estimated how much ATP tissues and tumors derived from glycolysis (2 ATP produced per glucose consumed) versus from TCA paired with the electron transport chain (approximately 14.5 ATP produced per two-carbon unit consumed). Tumors derived a much higher fraction of their ATP from glycolysis than healthy tissues, with an estimated 11-30% of ATP derived from glycolysis in tumors but a median of 1.8% derived from glycolysis in healthy tissues (Figure 5E).

We further used our measured glucose and TCA flux values to calculate total ATP production in healthy tissues and tumors. Remarkably, the implied ATP production rate of tumors was much less than healthy tissues. This calculation assumes a similar efficiency of ATP produced per reducing equivalent across tissues and tumors. Our calculated rates of ATP production are similar to studies measuring ATP phosphorylation rate in humans: around 6 μmol /min/g tissue in resting human soleus versus our measurement of 28 μmol /min/g tissue in working mouse diaphragm^44^. Tumors produced an average of 800 μmol ATP/minute/gram tissue, compared to an average of 10600 μmol ATP/minute/gram tissue in healthy tissues (Figure 5F-H). This finding stands in contrast to the widely held belief that cancer has a higher energy demand than healthy tissues.

How might tumors proliferate while using so much less ATP than healthy tissues? Healthy tissues use energy to carry out their physiological functions: the kidney pumps ions, the pancreas synthesizes digestive enzymes, et cetera. We hypothesize that tumors downregulate such physiological tasks, thereby conserving energy. To explore this possibility in the context of pancreatic cancer, we measured protein synthesis flux in pancreas and pancreatic cancer. Mice were infused with [U-^13^C] valine and its accumulation in total protein measured in pancreas and flank PDAC (Figure 5I). Indeed, pancreatic cancer carried out around four-fold less protein synthesis than the pancreas. Thus, at least in pancreatic tumors, downregulation of ATP production occurs in concert with loss of the native tissue’s hallmark ATP-consuming process.

## Discussion

Our kinetic tracing method revealed that the five tumor models surveyed have suppressed TCA cycle flux compared to healthy tissues. As expected, the tumors showed upregulated glycolysis, but nevertheless made most ATP oxidatively. Moreover, elevated glycolysis was insufficient to compensate for decreased TCA turning, with calculated ATP production rate significantly slower in the tumors than healthy tissues.

Accelerated glucose catabolism is the longest standing metabolic hallmark of tumors^9–11^, and supports the demands of proliferation. The consistent upregulation of glycolysis in tumors may in part reflect ATP demand not being satisfied by TCA flux, but also involves oncogene-driven metabolic rewiring, including upregulation of glucose transporters and glycolytic enzymes. This is mediated in part by activating mutations of the insulin signaling pathway (PI3 kinase pathway), which drive the Warburg effect^45,46^.

Upregulated glycolysis is, however, apparently not part of a general hypermetabolic program in tumors. While quiescent lymphocytes may need to broadly upregulate their metabolism to proliferate in response to an infection^46–48^, our data argue for a different metabolic strategy in solid tumors. Such tumors do not typically originate from cell types that are metabolically dormant like quiescent lymphocytes. Instead, they frequently derive from cells with substantial metabolic activity, including some of the most metabolic ones in the body like renal tubular epithelium. As such, these cells do not need to broadly ramp up all aspects of metabolism in order to proliferate. Instead, as they undergo oncogene-driven dedifferentiation, they lose energetically demanding tissue-specific functions such as digestive enzyme synthesis in the exocrine pancreas or ion pumping in the kidney^49,50^. This allows cancer cells to proliferate with decreased, rather than elevated, ATP production flux.

Data on mitochondrial membrane potential in tumors is consistent with limited ATP consumption sometimes gating TCA turning. Specifically, some lung cancers show elevated mitochondrial membrane potential^51^. A logical cause of such elevation is bottlenecking of ATP synthase activity due to limited ATP consumption, leading to proton accumulation in the mitochondrial inner membrane space. Thus, rather than tumors having defective mitochondria as Warburg hypothesized, they may sometimes have supercharged mitochondria with nowhere to put the available energy.

That said, cancer cells often face a hostile metabolic microenvironment where oxygen and nutrients are scarce, and competition with other cell types fierce^52–54^. The nefarious ability of cancer to proliferate with suppressed TCA turning and ATP demands positions them unfortunately well to succeed in the face of these metabolic challenges.

## Author Contributions

This work was conceived by C.R.B. and J.D.R. Y.S. helped advise on methods and quantative analysis. X. X. developed isocorr package which was used in analysis of mass spectrometry data. Experiments were carried out by C.R.B., W.D.L., T.T., C.S.R.J., L.W., L.Y., A.R.., V. B., T. L., Z.H., W.L., and J.Y.G.

## Acknowledgments

This work was funded by R01CA163591 from the National Cancer Institute to J.D.R., DP1DK113643 from the National Institute of Diabetes, Digestion and Kidney Disease to J.D.R., Ludwig Cancer Research funding to J.D.R., R50CA211437 from the National Cancer Institute to W.L., and Damon Runyon Postdoctoral Fellowship to C.R.B.

## Methods

### Mouse strains

All animal studies were approved either by the Princeton Institutional Animal Care and Use Committee (majority of experiments) or the Rutgers Institutional Animal Care and Use Committee (experiments with GEMM PDAC and GEMM NSCLC mice). Experiments in non-cancer bearing mice were performed in 9-14 week old C57Bl/6N mice from Charles River Laboratory. Mice were fed a standard rodent diet (PicoLab Rodent 5053 Laboratory Diet). For experiments with spontaneous pancreatic adenocarcinoma, Pdx1-cre;LSL-Kras-G12D/+;Trp53^fl/fl^ (‘GEMM PDAC’) mice were used at 5-8 weeks of age. For experiments with spontaneous lung adenocarcinoma, LSL-Kras-G12D/+; Trp53^fl/fl^; LKB1^fl/fl^ (‘GEMM NSCLC’) mice were inoculated intranasally with 4×10^7^ particles of Cre-expressing adenovirus; experiments were performed around 8 weeks after inoculation at 13-16 weeks of age.

### CRC xeno (HCT116) tumors

HCT116 colon cancer derived cells were grown in DMEM with 10% fetal bovine serum. For tumor implantation, cells were grown to confluency, then trypsinized and resuspended at 50×10^6^ per mL in media. Cells were mixed with Matrigel (Corning 354234) at a 1:1 (v/v) ratio, then injecting 200 microliters subcutaneously into the flank of CD1 nude mice (Charles River Laboratory strain 086) using 26G needle. Tumor growth was monitored by measuring the tumor dimensions (length, width and height) twice per week using a caliper. Tumor volume was calculated as 0.5 x (Length x Width x Height). Infusions were carried out 20-30 days after tumor implantation.

### Flank NSCLC tumors

Cell line was established as described in Davidson et al.^55^ from LSL-Kras^G12D/+^; Trp53^fl/fl^ mice inoculated intranasally with Cre-expressing adenovirus. Cells were grown in DMEM with 10% fetal bovine serum. For tumor implantation, cells were grown to 70% confluency, then trypsinized and resuspended at 2×10^6^ per mL in PBS, then 100 microliters was injected subcutaneously into the flank of C57/Bl6 mice. Tumor growth was monitored by measuring the tumor dimensions (length, width and height) twice per week using a caliper. Tumor volume was calculated as 0.5 x length x width x height). Infusions were carried out 25-35 days after tumor implantation.

### Flank PDAC tumors

Syngeneic pancreatic adenocarcinoma allograft tumors were established by harvesting tumors from Pdx1-cre;LSL-Kras-G12D/+; LSL-Trp53-R172H/+ mice, mincing the tissue into small particles, suspended in DMEM medium, mixing with Matrigel (Corning 354234) at a 1:1 (v/v) ratio, then injecting 200 microliters subcutaneously into the mouse flank. Allograft tumors were passaged up to two times in C57Bl/6 syngeneic recipient mice before implantation for use in experiments. Tumor growth was monitored by measuring the tumor dimensions (length, width and height) twice per week using a caliper. Tumor volume was calculated as 0.5 x (Length x Width x Height).

### Jugular vein catheterization

Aseptic surgical techniques were used to place a catheter (Instech C20PV-MJV1301 2Fr 10cm) in the right jugular vein and to connect the catheter to a vascular access button (Instech VABM1B/25 25gauge one-channel button) implanted under the back skin of the mouse. Mice were allowed to recover from surgery for at least 5 days before tracer infusion.

### Lactate primed infusion to measure TCA flux

[U-^13^C] lactate tracer (98% 13C, 20 w/w% solution, CLM-1579, Cambridge Isotope Laboratory) was diluted to 1.67% (3-fold more) in sterile water (not saline, to match osmolarity of blood). This tracer was stored at 4C until day of experiment. Jugular vein catheterized mice were fasted by switching to a fresh cage with no food at 9am. Around 1pm mice were connected to infusion line with swivel and tether (Instech products: swivel SMCLA, line KVABM1T/25) and infusion pump (syringepump.com, NE-1000), with infusate advanced through the tubing to the point of connection with the mouse. Mouse was left in cage connected to line for one hour to reduce stress. Around 2:30-3:30pm, infusion was initiated: prime dose of 13 microliters (to clear mouse catheter up to jugular vein)+0.48 microliters*(mouse weight in grams) was provided in 30 seconds, then infusion rate was slowed to 0.3 microliters*(mouse weight in grams) per minute. At the desired timepoint, mice were euthanized quickly by cervical dislocation, and tissues were collected as quickly as possible and freeze-clamped using a liquid nitrogen cooled Wollenberger clamp. Tissues were stored at −80C until processed.

### Glutamine primed infusion to measure TCA flux

[U-^13^C] glutamine (99% purity, CLM-1822) was diluted to 100mM in sterile saline. This tracer was made fresh for every experiment due to previously observed tracer instability. Infusion was performed as in previous section, with these modifications: priming dose of 13+1.6*(mouse weight) microliters was provided in 60 seconds, then infusion rate was slowed to 0.1 microliters*(mouse weight in grams) per minute. As above, at the desired timepoint, mice were euthanized quickly by cervical dislocation, and tissues were collected as quickly as possible and freeze-clamped using a liquid nitrogen cooled clamp. Tissues were stored at −80C until processed.

### 2-deoxyglucose primed infusion to measure glucose usage flux

[1-^13^C] 2-deoxyglucose (99% purity, CLM-1824) was diluted to the specified concentration (in most experiments, 8.3mM) in sterile saline. This tracer was stored at 4C until day of experiment. Infusion was performed as in previous section, with these modifications: priming dose of 13 microliters (to clear mouse catheter up to jugular vein) was provided in 30 seconds, then infusion rate was slowed to 0.3 microliters*(mouse weight in grams) per minute. As above, at the desired timepoint, mice were euthanized quickly by cervical dislocation, and tissues were collected as quickly as possible and freeze-clamped using a liquid nitrogen cooled clamp. Tissues were stored at −80C until processed.

### [U-^13^C] valine infusion to measure protein synthesis flux

[U-^13^C] valine (99% purity, CLM-2249) was diluted to 20mM in sterile saline. This tracer was stored at 4C until day of experiment. Infusion was performed as in previous section, with these modifications: infusion of 0.1 microliters*(mouse weight in grams) per minute for 3 or 4 hours (no priming dose).

As above, at the desired timepoint, mice were euthanized quickly by cervical dislocation, and tissues were collected as quickly as possible and freeze-clamped using a liquid nitrogen cooled clamp. Tissues were stored at −80C until processed.

### Blood enrichment sampling in double catheterized mice

Infusions were performed as above, but in mice purchased from Charles River Laboratory with carotid artery and jugular vein catheters. In that case the infusion line used was for double catheterized mice (Instech products: swivel SMCLA, line KVABM1T/25, second line and double-channel button VABM2T/25GCY, 25G tubing VAHBPU-T25, PNP3F25R pinport for sampling from arterial catheter).

Blood was sampled from arterial catheter at timepoints after the start of infusion using capillary blood collection tube (Sarstedt 16.440.100) and pinport-to-tube connector (Instech PNP3MS). Before and after sampling, arterial catheter line was flushed using a syringe and syringe-to-pinport connector (Instech PNP3M) with heparin-saline (1:50 heparin solution made with heparin SAI HGS-10). Blood was placed on ice, then centrifuged at 4C 14000 RCF 10min after all samples were collected. Serum was moved to another tube and stored at −80C.

For blood glucose concentration measurement, the same fasting and sampling procedure was performed but without any infusion.

### [U-^13^C] glucose infusion to measure whole-body glucose use

[U-^13^C] glucose (99% purity, CLM-1396) was diluted to 200mM in sterile saline. This tracer was stored at 4C until day of experiment. Infusion was performed using double catheterized mice as described above, with these modifications: infusion of 0.1 microliters*(mouse weight in grams) per minute for 2.5 hours (no priming dose).

### Blood and tissue processing and extraction

Throughout processing until extraction, tissues were kept on dry ice. Tissues were ground into powder using the Retsch CryoMill. Tissue powder was weighed (5-20mg in precooled Eppendorf tubes), and tissues were extracted by vortexing in 40x volumes precooled acetonitrile-methanol-water (40%/40%/20% v/v/v), then left on water ice over dry ice for 10 minutes. Then solution was centrifuged for 25min at 14000 RCF at 4C, moved to a new tube, then centrifuged again for 25min at 14000 RCF at 4C to remove any particulates. Then extract was moved to mass spectrometry vials (Thermo Scientific 200046, caps Thermo Scientific 501313) for measurement.

For blood samples, blood was kept on ice for up to 60min after sampling, then centrifuged at 4C 15000 RCF for 10 minutes. Serum fraction was transferred to another tube and stored at −80C. For mass spectrometry, 2-3 microliters of sample was extracted in 50 volumes methanol, then centrifuged at 4C 15000 RCF for 20 minutes, then transferred to a mass spectrometry vial for measurement.

To measure 2-deoxyglucose concentration in blood, samples were phosphorylated using purified hexokinase (Sigma H4502) according to the protocol described in Chiles et al.^56^ [U-^13^C] 2-deoxyglucose (Omicron Biochemicals, GLC-107) was phosphorylated using the same approach, 300uM in reaction mixture. Serum and standard (5-fold diluted to 60uM) were mixed and extracted in methanol (3ul serum+ 3ul standard+144 ul methanol).

To measure 2-deoxyglucose-phosphate concentration in tissues, a known concentration of [U-^13^C] 2-deoxyglucose (Omicron Biochemicals, GLC-107) was phosphorylated using the protocol above (300uM 2-deoxyglucose in the reaction mixture). This reaction mixture was then frozen and used as a standard when extracting tissues: standard was mixed into extraction buffer at a concentration of 0.25 micromolar (i.e. 10uM relative to tissues, which were extracted in 40x volume extraction buffer).

To measure TCA metabolite concentration in tissues, standards were made by resuspending ^13^C- or ^15^N-labeled metabolites (Cambridge Isotope Laboratories products: [U-^15^N] glutamate and [U-^15^N] aspartate from algal amino acid mix NLM-2161 [U-^13^C] a-ketoglutarate:CLM-4442, [U-^13^C] citrate:CLM-9021, [U-^13^C] malate: CLM-8065, [U-^13^C] succinate: CLM-1571) in water (at concentrations of 3040uM glutamate, 1980uM aspartate, 515uM a-ketoglutarate, 2693uM citrate, 483uM malate, 1365uM succinate). Standards were mixed into 40:40:20 (acetonitrile:methanol:water) extraction buffer at 1:100 concentration, and tissue powder samples were extracted in 100x volume extraction buffer on dry ice over water ice, then proceeded as above.

To measure glucose concentration in arterial blood, standards were made by resuspending ^13^C-labeled glucose (Cambridge Isotope Laboratories CLM-1396) in water at a known concentration, mixing standard into methanol extraction buffer, and blood serum samples were extracted in extraction buffer on water ice over dry ice, then proceeded as above.

To measure protein enrichment from [U-^13^C] valine infusion, tissues were ground using the Retsch Cryo Mill, then 5mg of tissue powder was weighed into Eppendorf tubes. Free tissue amino acids were removed using methanol-chloroform extraction as follows: 400ul methanol, 200ul chloroform, and 300ul water were added to each Eppendorf, vortexing after each addition. Tubes were centrifuged 10min at 14000 RCF at 4C, then upper layer was discarded. Remaining samples was washed twice with 500ul methanol, centrifuging 10min at 14000 RCF 4C between each wash. Finally, supernatant was removed and pellet was air dried. 250 ul 6M hydrochloric acid was added and tubes were incubated overnight at 98C. 10ul was moved to a new tube, then dried under nitrogen gas, resuspended in 500ul methanol and measured by mass spectrometry. (Note that each [U-^13^C] valine-infused tissue sample was also measured without methanol-chloroform extraction using the conventional sample-preparation method to measure free valine enrichment in the tissue.)

### Mass Spectrometry

Water soluble metabolite measurements were obtained by running samples on the Q Exactive Plus hybrid quadrupole-orbitrap mass spectrometer (Thermo Scientific) coupled with hydrophilic interaction chromatography (HILIC). An XBridge BEH Amide column (150 mm × 2.1 mm, 2.5 μM particle size, Waters, Milford, MA) was used. The gradient was solvent A (95%:5% H2O:acetonitrile with 20 mM ammonium acetate, 20 mM ammonium hydroxide, pH 9.4) and solvent B (100% acetonitrile) 0 min, 90% B; 2 min, 90% B; 3 min, 75%; 7 min, 75% B; 8 min, 70% B, 9 min, 70% B; 10 min, 50% B; 12 min, 50% B; 13 min, 25% B; 14 min, 25% B; 16 min, 0% B, 20.5 min, 0% B; 21 min, 90% B; 25 min, 90% B. The flow rate was 150 μL/min with an injection volume of 10 μL and a column temperature of 25°C. The MS scans were in negative ion mode with a resolution of 140,000 at m/z 200. The automatic gain control (AGC) target was 1 × 10^6^ and the scan range was m/z 75-1000.

In the case of 2-deoxyglucose-phosphate measurement, the method above was supplemented with a SIM scan to boost signal to noise ratio (minutes 11-15 of method, 240-255 m/z range).

All data were analyzed by El-MAVEN. For experiments involving carbon-13 labeling only (no nitrogen-15), outputs were corrected for natural ^13^C abundance using accucor package in R57.

For measuring concentrations of TCA intermediates in tissues, we used a mix of carbon-13 (13C) and nitrogen-15 (15N) labeled standards. We developed a software package in MATLAB to perform and visualize natural isotope correction for mass spectrometry data labeled with a mix of 13C and 15N (see https://github.com/xxing9703/MIDview_isocorrCN). This package requires that the instrumental resolution is high enough to fully resolve all the detectable 13C and 15N labeled peaks of a given compound, with a total number of possible peaks n:

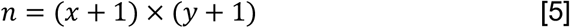

where x and y represent the number of carbon atoms and of nitrogen atoms in the compound respectively. Briefly, a correction matrix A of size n by n is constructed to relate the measured isotope labeling pattern before correction (L) to the isotope labeling pattern after correction (L_C_):

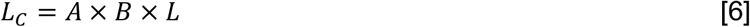

Each matrix element A_ij_ represents the jth labeled fraction contributing to the ith measured mass fraction. The n by n impurity matrix B is constructed to take into account impurities in 13C and 15N tracers (a value reported by the tracer manufacturer: typically 99% purity is used). Finally, we solve for L_C_ by non-negative least square fitting using the lsqnonneg solver in MATLAB.

### MALDI imaging mass spectrometry

Primed infusion of [U-^13^C] lactate was performed as described above for 90 seconds, or steady lactate infusion was performed for 150 minutes to measure steady-state labeling of TCA metabolites from lactate. Kidneys were quickly dissected and snap frozen on dry ice, then stored intact at −80°C until MALDI-IMS analysis. For each tissue, ~10 μm-thick sections were collected on a cryostat (Leica CM3050S, Wetzlar, Germany). Sections for MALDI-MSI were thaw-mounted on indium tin oxide (ITO)-coated glass slides (Bruker Daltonics, Bremen, Germany) and desiccated under vacuum for 10 mins. Alternating serial sections for immunohistochemistry and immunofluorescence were collected on standard glass slides.

Desiccated tissue sections mounted on ITO glass slides were sprayed using an HTX TM-Sprayer (HTX Technologies, LLC) with 10 mg/ml N-(1-Naphthyl) ethylenediamine dihydrochloride (NEDC, Sigma #222488) dissolved in 70%:30% methanol:water. The sprayer temperature was set to 80°C, with a flow rate of 0.1 ml/min, velocity of 1000 mm/min, track spacing of 2 mm, pressure of 10 psi, and 3 liters/min gas flow rate. Ten passes of the matrix were applied to slides with 10 seconds of drying time between each pass.

For MALDI-FTICR measurements, the matrix-coated slides were immediately loaded into a slide adapter (Bruker Daltonics, Bremen, Germany) and then into a solariX XR FTICR mass spectrometer equipped with a 9.4T magnet (Bruker Daltonics, Bremen, Germany). The resolving power 120,000 at m/z 500. Mass accuracy was calibrated within 1 ppm by using 1mg/mL Arginine solution before starting the run. m/z 124.0068 (taurine), m/z 133.0136 (malate) and m/z 145.0611 (glutamate) were used as lock masses during the run because of their high abundance in kidney. The laser focus was set to ‘minimum,’ and the x-y raster width was set to 20 μm using Smartbeam-II laser optics. A spectrum was accumulated from 200 laser shots at 1200 Hz, and the ions were accumulated using the “cumulative accumulation of selected ions mode” (CASI) within an m/z range of 70-250 before being transferred to the ICR cell for a single scan.

### Immunofluorescence

At time of harvesting, tumor samples were snap-frozen on dry ice for 30 min and stored at −80°C until sectioning. 10 μm-thick tumor sections were collected on a cryomicrotome (Leica CM3050S, Wetzlar, Germany) and mounted on slides. Slides were brought to room temperature and fixed with 4% paraformaldehyde, then washed with PBS and then PBS+0.1%Triton. Slides were stained using Lotus Tetragonolobus Lectin-fluorescein (Vector Laboratories FL-1321) in combination with M.O.M. Immunodetection Kit (Vector Laboratories BMK-2202) using the manufacturer’s protocol. Slides were counterstained with DAPI and mounted using Fluoromount-F (Thermo Fisher 00-4958-02), then imaged using a Cytation 5 microscope.

### Calculations and Analysis

See Extended Data Table 1 for a list of sample sizes for all experiments. Each sample is an independent biological replicate (either one tissue from an endpoint experiment, or a blood timepoint).

### TCA flux calculations

For all calculation of TCA cycle labeling, timepoints from 0 to 90 or 0 to 150 minutes were used: at least five timepoints per tissue, and for the majority of tissues, at least 12 timepoints per tissue. For carbon-13 lactate infusion, m+2 labeling of glutamate, malate, and succinate was averaged for each sample, while for carbon-13 glutamine infusion, m+5 labeling, m+4 succinate, and m+4 malate labeling was averaged for each sample. K labeling constants were calculated using nonlinear fitting of a single exponential-decay function using the R nls() function. To calculate TCA pool size, concentrations of glutamate, malate, succinate, aspartate, a-ketoglutarate, and citrate (calculated by internal standard spikein as described above) were summed for each organ. TCA flux was calculated as k labeling constant multiplied by TCA pool size. For the case of TCA flux measurement by glutamine infusion in pancreatic cancer models, since we had sampled fewer timepoints, we estimated the asymptote (eventual steady state labeling) as the mean of the 150min timepoint and fit a single-exponent curve to the 5min timepoint data. Note that TCA flux in brain could not be measured using either [U-^13^C] lactate or glutamine, since neither of this feeds the TCA cycle in brain substantially^25^.

In using this approach to calculate TCA flux, several assumptions were made. TCA metabolites in a tissue must be well-mixed and not compartmentalized. Precursors to TCA (e.g. serum lactate and tissue lactate) should label much faster than TCA labeling, otherwise their labeling speed might influence the calculated TCA flux.

To calculate oxygen consumption from TCA flux, we used Equation [3]:

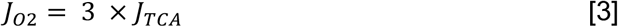

When lactate is oxidized, it yields 5 NADH (from the lactate dehydrogenase, pyruvate dehydrogenase, isocitrate dehydrogenase, alpha-ketoglutarate dehydrogenase, and malate dehydrogenase reactions) and 1 FADH2 (from the succinate dehydrogenase reaction). When a two-carbon unit of fatty acid is oxidized, it yields 4 NADH (from the fatty acid oxidation, isocitrate dehydrogenase, alpha-ketoglutarate dehydrogenase, and malate dehydrogenase reactions) and 2 FADH2 (from the fatty acid oxidation and succinate dehydrogenase reactions). Therefore each nutrient contributes 6 reducing equivalents per TCA turn (per acetyl-CoA consumed). Each reducing equivalent results in consumption of ½ O_2_ molecule in the electron transport chain, so each TCA turn leads to 3 O_2_ molecules consumed.

### Glucose usage calculation

To calculate glucose usage flux, 2-deoxyglucose-phosphate concentration in tissues was determined at timepoints from 0 to 15 minutes. This equation assumes that 2-deoxyglucose is infused in trace amounts that do not inhibit endogenous glucose metabolism, and indeed the peak 2-deoxyglucose concentration measured in blood was ~90 micromolar, or around 86-fold lower than blood glucose concentration (Figure 3B). We found a linear relationship between the rate of 2-deoxyglucose infused and the tissue 2-deoxyglucose-phosphate concentration (Extended Data Figure 3D) again suggesting the trace amount of 2-deoxyglucose infused was not saturating tissue glucose use. Each healthy tissue was measured from at least n=2 mice at 5min, 7.5min, 10min, 15min, and one 0min timepoint; tumors were measured from at least n=2 mice at 5min, 10min, and 15min except for GEMM NSCLC, which included only n=1 for 5min. 2-deoxyglucose concentration in blood was measured in arterial blood using double catheterized mice, and integrated concentration over time was calculated. The slope of the 2-deoxyglucose-phosphate tissue concentration versus blood 2-deoxyglucose integrated concentration curve was calculated using linear fitting in R. The glucose usage flux for each tissue was calculated as the slope of this line multiplied by the blood glucose concentration (7.7mM in mice fasted from 9am-3:30pm, Extended Data Figure 3C).

Note that glucose usage flux is similar to glycolysis flux, except that glucose usage flux measures glycolysis plus glycogen synthesis flux from glucose, while glycolysis includes glycolysis from both glucose and glycogen; quantitatively, the values are likely to be very similar in the fasted state since not much glycogen is synthesized.

Two tissues were analyzed differently: brown adipose demonstrated saturation of 2-deoxyglucose phosphate after 10 minutes, so the 15-minute timepoint was not used in further analysis (Extended Data Figure 3B). Colon 2-deoxyglucose phosphate displayed potential signal to noise problems that we are exploring further, so colon glucose usage rate was not analyzed.

### Whole body metabolism calculations

To calculate contribution of each tissue to whole-body TCA flux or glucose flux, the measured flux was multiplied by the fraction of body mass made up of each tissue, with tissue fractional masses from previous studies^30–33^. We had to estimate what fraction of total muscle consisted of what we call ‘white muscle’ (similar to quadriceps and gastrocnemius) versus red muscle (similar to diaphragm and soleus). Muscle fibers are classified into types I, IIA, and IIB, where I is ‘reddest’, IIA is intermediate, and IIB is ‘white’. Considering the mass of each measured muscle and its composition, we estimated that total mouse muscle is 3% type I fiber, 41% type IIA, and 50% type IIB. Quadriceps muscle and gastrocnemius muscle (what we are calling ‘white’) is 0% type I, 30% type IIA, 70% type IIB. The mean of diaphragm and soleus (what we are calling ‘red’) composition is 25% type I, 42% type IIA, 27% type IIB. Therefore, we used least squares fitting to this linear system of equations to estimate that mouse muscle is 41% red muscle and 59% white muscle. Since muscle in total makes up ~38% of the mouse body^30^, this means red muscle is approximately 16% and white muscle is approximately 23% of whole body mass. Red muscle TCA and glucose usage flux were estimated as the mean of diaphragm and soleus fluxes. White muscle TCA and glucose usage flux were estimated using the means of quadriceps and gastrocnemius fluxes.

### Lactate consumption/production calculation

To estimate whether a tissue consumed or released net lactate, we estimated net lactate production as 2 times the glucose usage flux from equation [4] (see equation [5]). We calculated net lactate consumption in the TCA cycle as the TCA flux from equations [1] and [2] by the contribution of lactate to the TCA cycle (see equation [6]) The contribution of lactate to the TCA cycle was the ratio of the mean value of glutamate, malate, and succinate m+2 enrichment at 90min of [U-^13^C] lactate primed infusion, divided by the mean value of arterial blood m+3 lactate enrichment from 45 seconds to 60 minutes (around 0.22).

### ATP production rate calculation

ATP production from glycolysis was calculated as 2 ATP per glucose consumed (we did not consider ATP derived from the NADH from the GAPDH reaction, as this is generally consumed by the lactate dehydrogenase reaction). ATP production from the TCA cycle was calculated thus: for lactate, 1 ATP equivalent (succinyl coA synthetase reaction), 5 NADH (lactate dehydrogenase, pyruvate dehydrogenase, isocitrate dehydrogenase, a-ketoglutarate dehydrogenase, and malate dehydrogenase reactions) and 1 FADH2 (succinate dehydrogenase reaction) are produced. From one 2-carbon unit of fatty acid, 1 ATP equivalent (succinyl coA synthetase reaction), 4 NADH (fatty acid oxidation, isocitrate dehydrogenase, a-ketoglutarate dehydrogenase, and malate dehydrogenase reactions) and two FADH2 (fatty acid oxidation and succinate dehydrogenase reaction) are produced. One NADH generates approximately 2.5 ATP in the electron transport chain while one FADH2 generates approximately 1.5 ATP. Thus for the two most common substrates, one TCA cycle turn generates either 14 or 15 ATP, so estimated 14.5 ATP per TCA turn.

### Protein synthesis rate calculation

To calculate protein synthesis rate, fractional enrichment of valine in protein was normalized to fractional enrichment of free valine in tissue from [U-^13^C] valine infusion. This fractional protein synthesis during the time of the infusion, which was 3-4 hours, was divided by the length of that infusion to yield fractional protein synthesis per hour.

#### Statistics

All t-tests used were two-sided; sample size for every experiment is recorded in Extended Data Table 1.

To compute two-sided t tests for difference in TCA fluxes, the following procedure was used: standard deviation of k labeling rate constant was computed by nls() function in R. Standard deviation of TCA pool size concentration was computed as the square root of the sum of squared standard deviations of the six TCA metabolite concentration measurements. The standard deviation of the TCA flux (the product of k labeling rate constant and TCA pool size concentration) was computed as the sum of the percent standard deviations: standard deviation of k divided by k, plus standard deviation of pool size divided by pool size, all multiplied by flux. The p value of a two-sided t test comparing two TCA fluxes was then computed using a Welch’s t test using the Welch-Satterthwaite equation (not assuming equal variances).

To compute two-sided t tests for difference in glycolysis fluxes, a similar procedure was used: standard deviation of the slope of the tissue 2-deoxyglucose-phosphate concentration versus the integrate serum 2-deoxyglucose concentration with respect to time was calculated by the lm() function in R. The standard deviation of the glycolysis flux (the product of the slope and the blood glucose concentration) was computed as the sum of the percent standard deviations: standard deviation of the slope divided by the slope, plus standard deviation of serum glucose concentration divided by mean serum glucose. The p value of a two-sided t test comparing two glycolysis fluxes was then computed using a Welch’s t test using the Welch-Satterthwaite equation (not assuming equal variances).

To compute two-sided t tests for difference in ATP production fluxes, the standard deviation was computed as the sum of squared standard deviations of glycolysis flux and TCA flux. The p value of a two-sided t test comparing two ATP production fluxes was then computed using a Welch’s t test using the Welch-Satterthwaite equation (not assuming equal variances).

All R^2^ values of linear correlation were computed using linear regression.

### Data Availability

All raw data, analyzed data and materials will be provided upon request to the corresponding author. We plan to also make the data available through an in-house data repository that is currently in development for easy retrieval and processing of stable isotope tracing mass spectrometry data.

