## Extended Data Figures for "Slow TCA flux implies low ATP production in tumors"

### Extended Data Figure 1

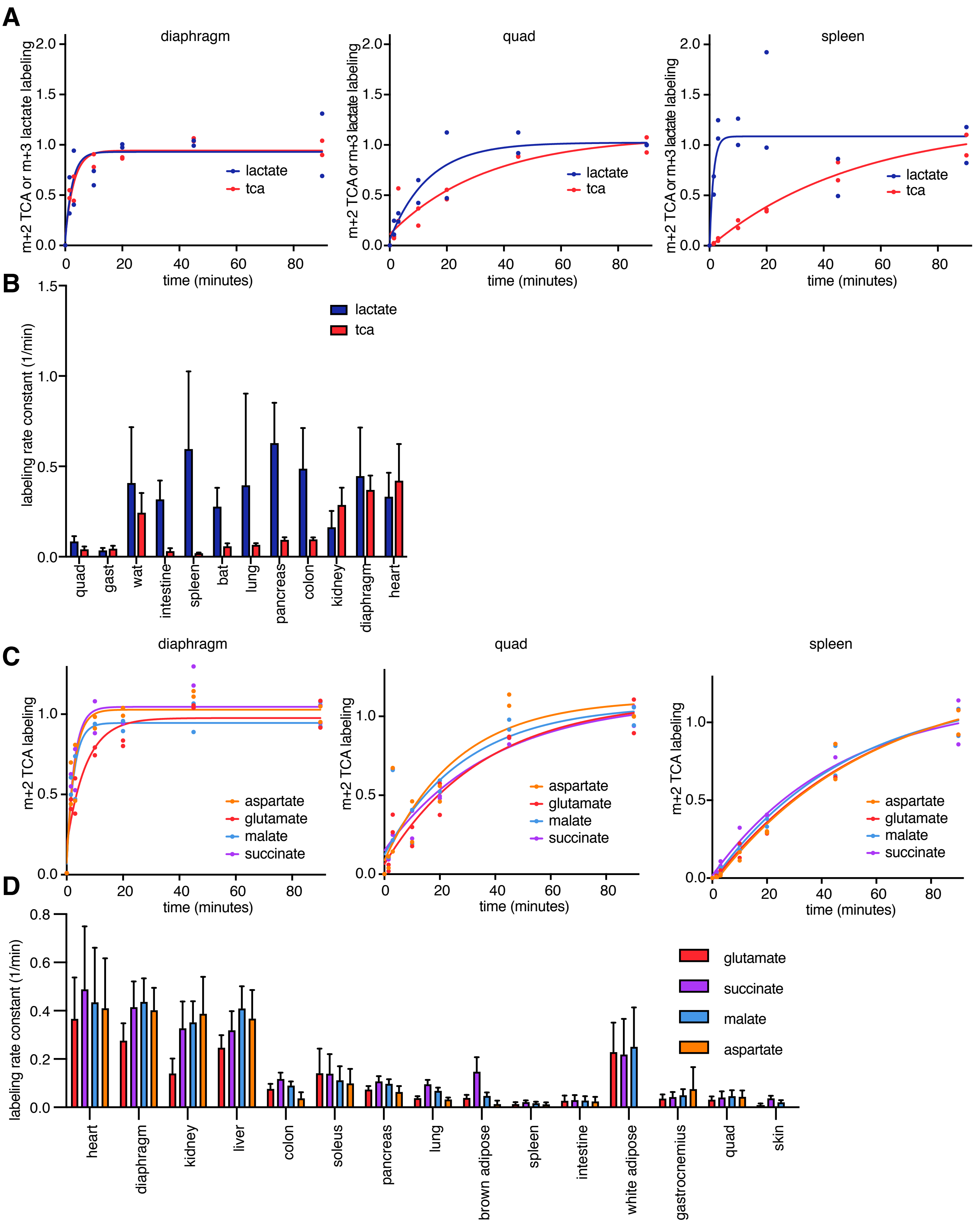


Extended Data Figure 1. TCA labeling from carbon-13 lactate primed infusion. (A) Tissue lactate m+3 and TCA m+2 (mean of glutamate, malate, succinate) labeling from carbon-13 lactate infusion. (B) Labeling rate constant of tissue lactate and TCA metabolites from carbon-13 lactate infusion. For accurate TCA flux measurement, tissue lactate labeling (blue bars) must be much faster than TCA labeling (red bars). (C) Tissue TCA intermediate m+2 labeling from carbon-13 lactate infusion. (D) Labeling rate constants of tissue lactate and TCA metabolites from carbon-13 lactate infusion. Metabolites in the same tissue label at similar rates, supporting the inclusion of glutamate and aspartate as part of the effective TCA metabolite pool.

### Extended Data Figure 2

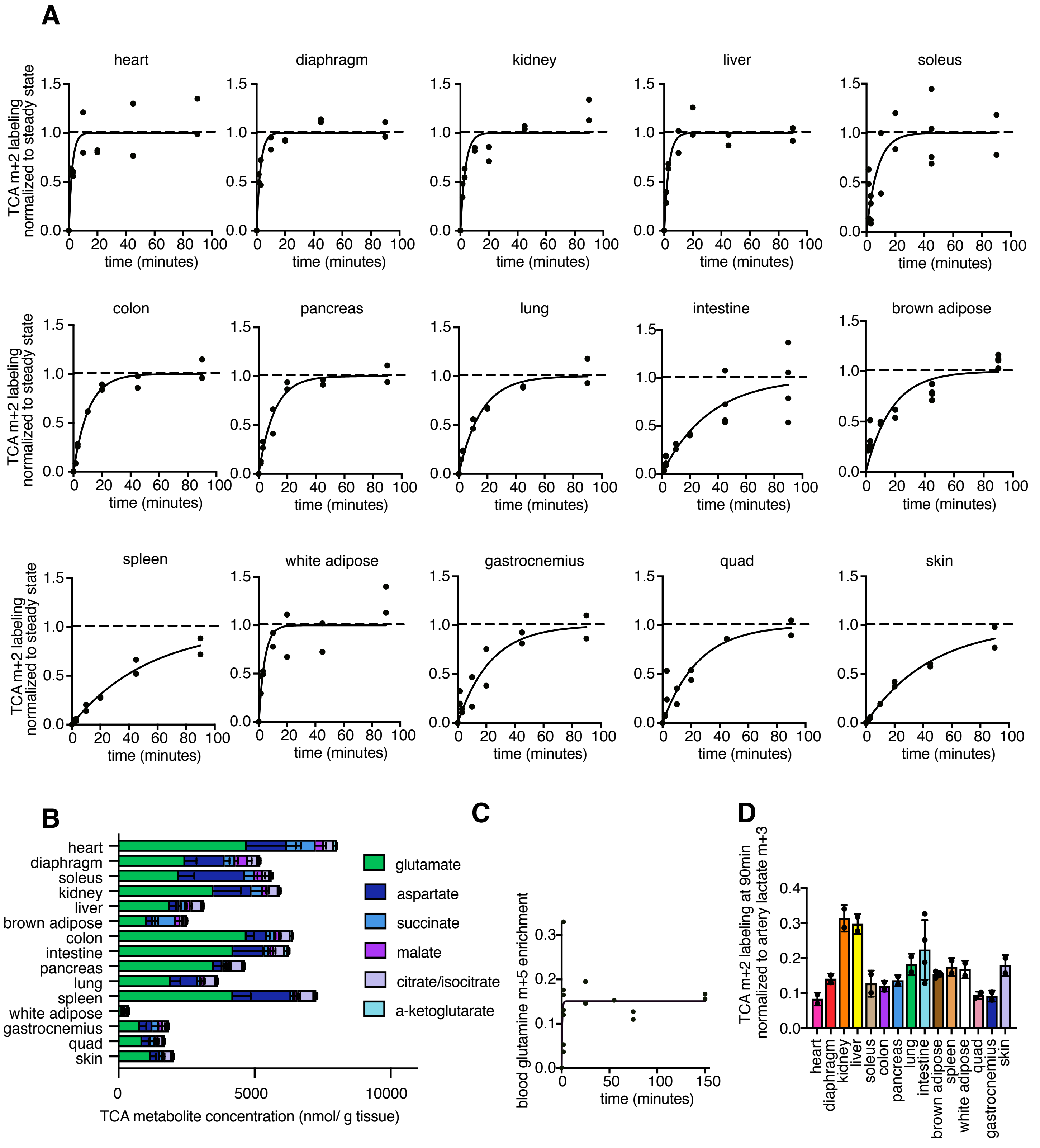


Extended Data Figure 2. TCA flux measurement. (A) TCA m+2 labeling timepoints from carbon-13 lactate primed infusion.
(B) Individual metabolites making up total TCA pool concentration. Error bars are standard deviation.
(C) Serum enrichment in carbon-13 glutamine primed infusion.
(D) Fractional contribution of lactate to tissue TCA. Error bars are standard deviation.

### Extended Data Figure 3

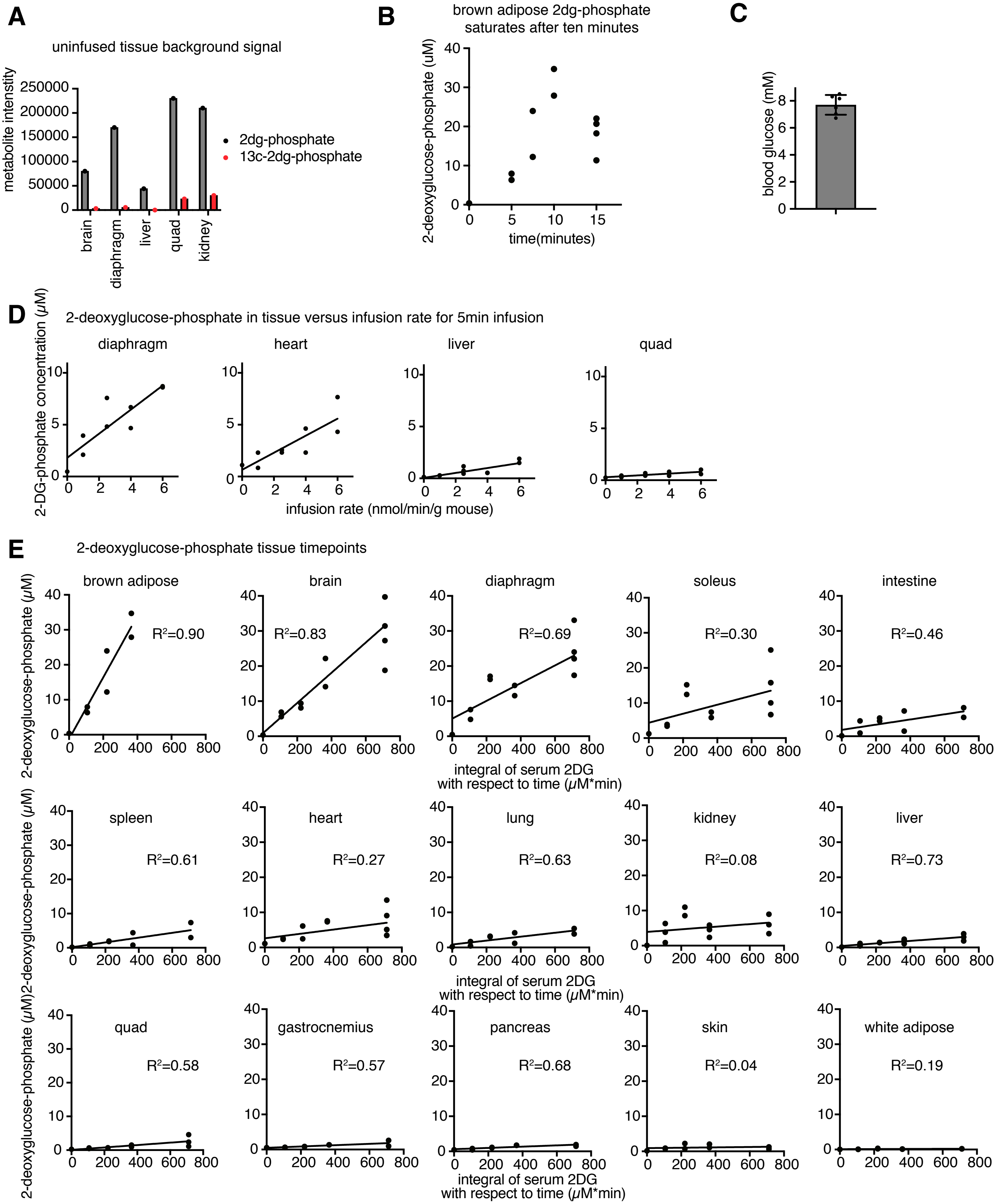


Extended Data Figure 3. Glucose usage flux measurement. (A) [1-$^{13}$C] 2-deoxyglucose phosphate has lower background signal in tissues compared to carbon-12 2-deoxyglucose phosphate. (B) Brown adipose 2-deoxyglucose tissue uptake and phosphorylation saturates after 10 minutes so later timepoint was discarded in calculating glucose use flux. (C) Arterial blood glucose concentration. (D) Tissue 2-deoxyglucose phosphate concentration versus 2-deoxyglucose infusion rate (5min timepoint). Linearity suggests that at 2.5nmol/min/g mouse infusion rate, 2-deoxyglucose uptake and phosphorylation is not saturated. (E) 2-deoxyglucose phosphate in tissues versus integral of serum 2-deoxyglucose over time (2.5nmol/min/g mouse infusion rate, 0-15min timepoints).

### Extended Data Figure 4

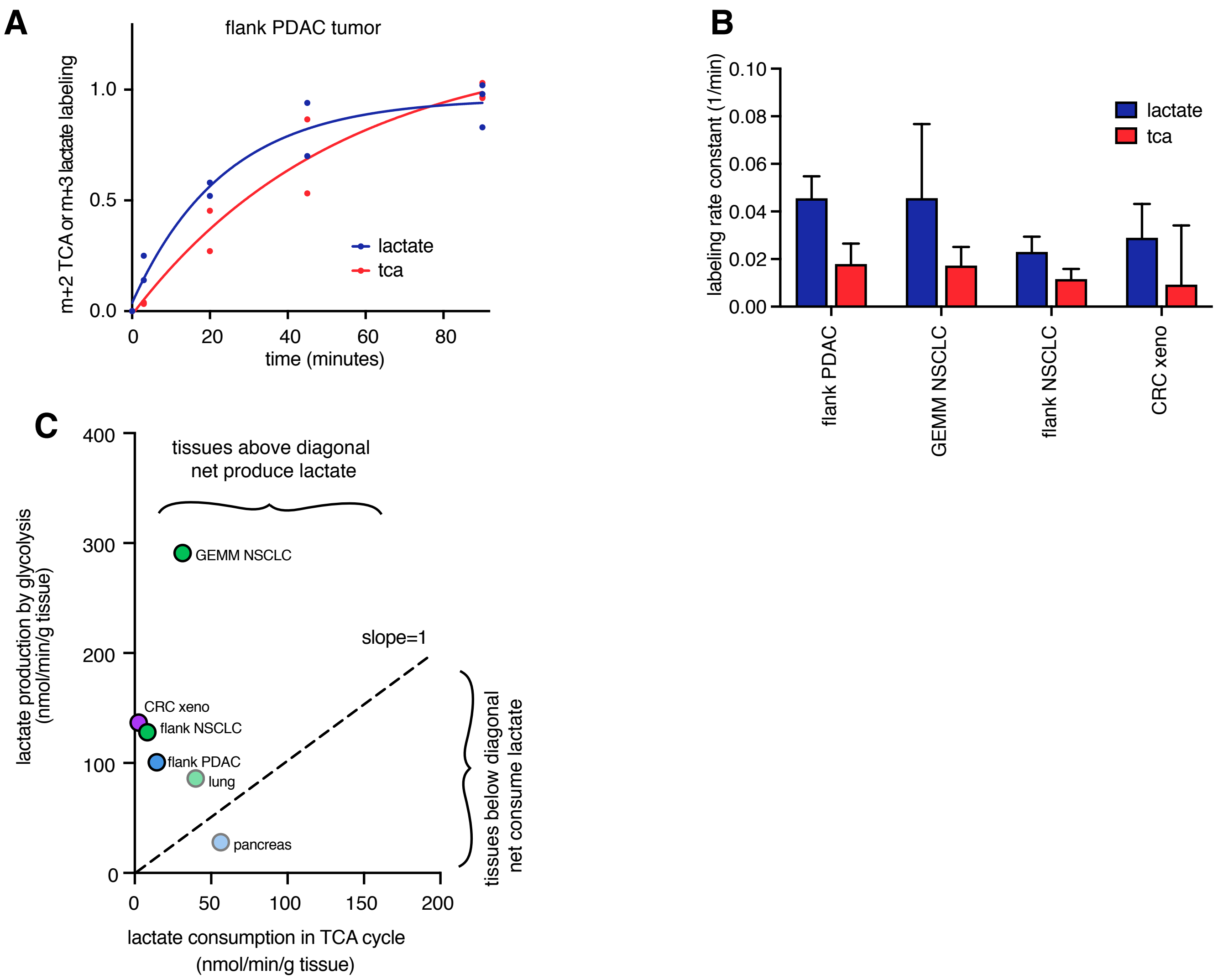


Extended Data Figure 4. Lactate and TCA labeling rate from carbon-13 lactate primed infusion. (A) Tissue lactate m+3 and TCA m+2 (mean of glutamate, malate, succinate) in flank PDAC tumors. (B) Labeling rate constants of tissue lactate and TCA metabolites in tumors from carbon-13 lactate infusion. (C) Production and consumption of lactate by healthy tissues and tumors calculated from glucose usage flux and from TCA flux measured by carbon-13 lactate multiplied by lactate contribution to TCA cycle

### Extended Data Table 1

| | blood | brain | brown adipose | colon | diaphragm | gastrocnemius | heart | intestine | kidney | liver | lung | pancreas | quad | skin | soleus | spleen | white adipose | GEMM PDAC | flank PDAC | GEMM NSCLC | flank NSCLC | xeno CRC |
|---|---|---|---|---|---|---|---|---|---|---|---|---|---|---|---|---|---|---|---|---|---|---|
| non-primed carbon-13 lactate infusion | 12 | | | | | | | | | | | | | | | | | | | | | |
| carbon-13 lactate primed infusion timepoints | 41 | | 19 | 12 | 12 | 12 | 12 | 19 | 12 | 12 | 12 | 12 | 12 | 11 | 17 | 12 | 12 | | 9 | 9 | 10 | 7 |
| tca pool size | | | 4 | 4 | 4 | 4 | 4 | 4 | 4 | 4 | 4 | 4 | 4 | 4 | 4 | 4 | 4 | 4 | 4 | 4 | 4 | 4 |
| carbon-13 glutamine primed infusion timepoints | 14 | | | | | | | 8 | 8 | 8 | 8 | 8 | 8 | | | 8 | | 5 | 10 | | | |
| lactate primed infusion+imaging mass spectrometry | | | | | | | | | 4 | | | | | | | | | | | | | |
| 2-deoxyglucose infusion (2.5nmol/min/g mouse) | 12 | 13 | 13 | 16 | 13 | 9 | 13 | 9 | 12 | 12 | 9 | 9 | 12 | 9 | 13 | 9 | 9 | | 6 | 5 | 6 | 6 |
| protein synthesis measurements (carbon-13 valine infusion) | | | | | | | | | | | | 6 | | | | | | | 4 | | | |
| blood glucose concentration measurement | 6 | | | | | | | | | | | | | | | | | | | | | |
| 2-deoxyglucose infusion (1nmol/min/g mouse, 5 minute timepoint) | | | | | 2 | | 2 | 2 | 2 | 2 | | | 2 | | | | | | | | | |
| 2-deoxyglucose infusion (4nmol/min/g mouse, 5 minute timepoint) | | | | | 2 | | 2 | 2 | 2 | 2 | | | 2 | | | | | | | | | |
| 2-deoxyglucose infusion (6nmol/min/g mouse, 5 minute timepoint) | | | | | 2 | | 2 | 2 | 2 | 2 | | | 2 | | | | | | | | | |

Extended Data Table 1: N (independent tissue or blood samples) for each experiment. Every sample was measured once.
